# Multiple parietal pathways are associated with rTMS-induced hippocampal network enhancement and episodic memory changes

**DOI:** 10.1101/2020.07.09.195172

**Authors:** Michael Freedberg, Catherine A. Cunningham, Cynthia M. Fioriti, Jorge Dorado Murillo, Jack A. Reeves, Paul A. Taylor, Joelle E. Sarlls, Eric M. Wassermann

## Abstract

Repetitive transcranial magnetic stimulation (rTMS) of the inferior parietal cortex (IPC) increases resting-state functional connectivity (rsFC) of the hippocampus with the precuneus and other posterior cortical areas and causes proportional improvement of episodic memory. The anatomical pathway responsible for the propagation of these effects from the IPC is unknown and may not be direct. Using diffusion tensor imaging, we examined whether individual differences in fractional anisotropy (FA), a tensor-derived quantity related to white matter properties, in pathways between the IPC and medial temporal lobe (MTL), via the parahippocampal cortex and the precuneus, accounted for individual differences in hippocampal rsFC and memory change after rTMS. FA in the IPC-parahippocampal pathway was associated with rsFC change in a few small cortical clusters, while FA in the IPC-precuneus pathway was strongly linked to widespread changes in rsFC. FA in both pathways was related to episodic memory, but not to procedural memory. These results implicate pathways to the MTL and to the precuneus in the enhancing effect of parietal rTMS on hippocampal rsFC and memory.

## Introduction

Repetitive transcranial magnetic stimulation (rTMS) can modulate resting-state functional connectivity (rsFC) in brain networks (Halko et al. 2014; Wang et al. 2014; Rastogi et al. 2017; Freedberg et al. 2019; Hermiller et al. 2019). By delivering rTMS to the inferior parietal cortex (IPC), Wang et al. (2014) increased rsFC between the hippocampus and a network of posterior cortical regions, most prominently the precuneus. The magnitude of this change was linearly related to improved episodic memory among individuals. These effects have been reproduced several times by the same (Hermiller et al. 2019; Warren et al. 2019) and other (Tambini et al. 2018; Freedberg et al. 2019) laboratories, and they appear to be specifically associated with rsFC increases in a set of cortical regions with high baseline hippocampal rsFC. Using rTMS to increase memory network connectivity may help patients with pathologies that cause reduced efficiency in networks involved in episodic memory (e.g. Alzheimer’s disease; Greicius et al. 2004)

Previous studies have targeted the IPC sub-region with the strongest hippocampal rsFC within an individual because the hippocampus is the critical hub of the episodic memory network and because there is a related assumption that an IPC-hippocampus connection is central to the effect (Wang et al. 2014; Freedberg et al. 2019; Hermiller et al. 2019; Warren et al. 2019; Freedberg et al. 2020). One proposed route for propagation of hippocampal rsFC-reinforcing activity from the stimulated site is via retrosplenial and/or parahippocampal cortex to the hippocampus, and thence, via divergent pathways, to regions connected to the hippocampus (Wang et al. 2014; Freedberg et al. 2019; Hebscher and Voss 2020). However, to date there is no experimental evidence for this hypothesis.

High rsFC between regions can occur via intermediate nodes (Honey et al. 2009; Warren et al. 2017) where anatomical connectivity exists (Gong et al. 2009; Davis et al. 2017), and models have been able to explain the effects of TMS on network rsFC based on anatomical connections (Muldoon et al. 2016). Individual differences in pathway anatomy also have the potential to explain the wide variability in the physiological (López-Alonso et al. 2014) and behavioral (Nicolo et al. 2015) responses to rTMS.

Diffusion tensor imaging (DTI) analysis of diffusion-weighted (DW) MRI can detect individual variations in white matter (WM) pathways due to its sensitivity to tissue microstructure. DTI modeling allows estimation of the location of major WM pathways (via tractography), as well as the quantification of their properties. Fractional anisotropy (FA) is a scalar value (Basser 1995; Jones et al. 2013) that can highlight important differences between individuals (Cheng et al. 2012) and is sensitive to axon diameter, fiber density, membrane permeability, myelination, and the intra-voxel orientational coherence of axons (Beaulieu 2002). FA has been used in conjunction with physiological methods (Honey et al. 2009; Betzel et al. 2014) to explain inter-individual rsFC variability.

In order to study the mechanism underlying the effect of IPC rTMS on hippocampal rsFC and episodic memory, we used DTI-based tractography to estimate the segments of the WM skeleton most likely connecting target-relevant regions of interest (ROIs). We then measured FA in pathways capable of transmitting the effect to the hippocampus and other nodes in the episodic memory network to determine locations with strong associations between individual anatomy and response.

We measured FA in two independent candidate pathways linking the stimulation point to downstream regions. The first connected the IPC to a region of the medial temporal lobe (MTL), including the parahippocampal and entorhinal areas. We did not posit a direct pathway from the IPC to the hippocampus because no such pathway has been found by anatomical tracing in primates or DTI in humans. Typically, tracing studies have focused on area 7 (IPC, or posterior parietal cortex) to link it to the parahippocampal cortex (Mesulam et al. 1977; Pandya and Seltzer 1982; Cavada and Goldman-Rakic 1989a). The parahippocampal cortex projects to the caudal entorhinal cortex (Suzuki and Amaral 1994) and the entorhinal cortex then projects to the hippocampus via the perforant pathway (Amaral and Witter 1989). Thus, we examined IPC-MTL connections in three segments: 1) IPC to parahippocampal cortex, 2) parahippocampal cortex to entorhinal cortex, and 3) entorhinal cortex to hippocampus.

The other pathway we examined connects the IPC to the precuneus, the region showing the most reproducible and robust hippocampal rsFC increase after IPC rTMS (Wang et al. 2014; Freedberg et al. 2019). Anatomical tracing studies in monkeys (Pandya and Seltzer 1982; Cavada and Goldman-Rakic 1989a; Morecraft et al. 2004) have shown projections from area 7 (IPC) to area 7m (precuneus) and from there to the MTL, and similar pathways have been shown with DTI in humans (Takahashi et al. 2008). The precuneus has diverse cognitive functions (Cavanna and Trimble 2006), but, importantly for us, it is critical for episodic memory (Valenstein et al. 1987; Rudge and Warrington 1991; von Cramon and Schuri 1992).

Enhancement of rsFC in a network functionally centered on the precuneus, rather than on the hippocampus, is also an alternative explanation for the effect of IPC rTMS on memory.

To investigate that the effects of IPC rTMS on behavior were specific to episodic memory, we also tested procedural memory, which involves a nominally separate system based principally in a network of frontal and striatal regions (Poldrack et al. 1999). We expected that there would be no association between FA in either of these pathways and rTMS-induced changes in procedural learning.

## Materials and Methods

### Subjects

Forty-eight adults (20 female; mean age = 25.33 ± 4.80 yrs), free of neurological or psychiatric disorders or medications acting on the central nervous system, participated in one of two studies. All participants passed screening for contraindications to TMS (Rossi et al. 2009) and MRI. Fifteen participated in a rTMS dose-finding study and received one, three, or four daily consecutive sessions of IPC stimulation, but not memory testing. The remaining 33 participated in a study examining the effect of IPC rTMS on hippocampal rsFC relative to vertex control stimulation. Eighteen of these participants received three daily consecutive sessions of IPC stimulation, while the remaining 15 received vertex stimulation on an identical schedule. All participants in this experiment received episodic and procedural memory testing. From these 48 participants, we excluded 13 for the following reasons: received only one session of stimulation (3); the hippocampal seed used to locate the IPC target was mistakenly placed in the parahippocampal gyrus and used to guide rTMS (3); failure to follow instructions during baseline behavior testing (1); excessive head motion during scanning (1); incomplete or technically poor DTI data acquisition (5). This left 35 participants (16 female; mean age = 25.31 ± 4.52 yrs). Of this group, 18 received three sessions of IPC stimulation, six received four sessions of IPC stimulation, and 11 received three sessions of vertex stimulation. Of the 35 participants, behavioral testing was included for 12 participants from the IPC stimulation group and 11 from the vertex group. One participant from the IPC group developed an explicit strategy during the baseline procedural memory test and we omitted their data for that test. All participants in the final data set reported being right-handed. Written informed consent was obtained and the study was approved by the local Institutional Review Board.

### Procedures

All subjects underwent, in order: baseline MRI scanning, three or four consecutive daily rTMS sessions, and a post-stimulation MRI scan. Behavioral measures were taken immediately following each scan and 7-14 days after the last rTMS session (Fig. 1A). Baseline MRI scanning included an anatomical localizer, structural scan (for functional scan co-localization with anatomy, and neuro-navigation), a resting state fMRI scan, and DW scans. Participants underwent their first rTMS session within 36 hours of baseline scanning. The second MRI session occurred on the day after the final rTMS session and within three hours of the time of day of the first scanning session. Subjects were blinded to the specific intent of the study and to the stimulation condition.

**Figure 1.**
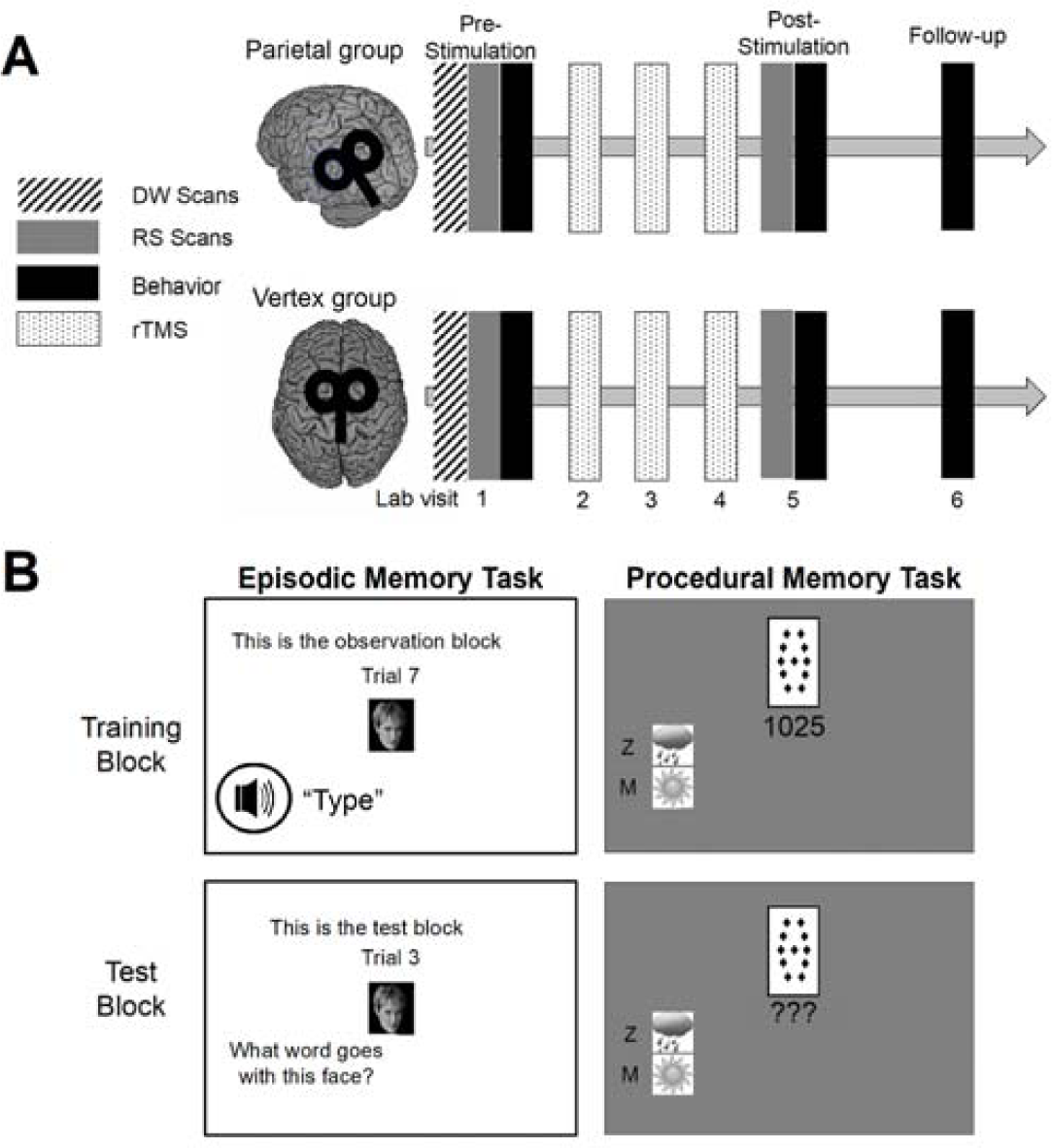
A. Timing of procedures and stimulation sessions. The first session of stimulation occurred within 36 hours of the initial assessments. Six of the participants received four sessions of stimulation. Follow-up testing occurred 7-14 days after the last rTMS session. DW = diffusion-weighted, RS = resting-state. B. Memory tasks. (left) Episodic memory task.

Participants encoded face-word pairs during the training block and needed to recall the paired word during testing. (right) Procedural memory task. During training, participants experienced 14 different arrangements of four different cards and needed to learn the probabilistic relationship between the cards and two possible outcomes through feedback. During testing, participants were tested on their ability to produce the optimal response for each arrangement.

### fMRI and DWI acquisition

MRI was performed on a Siemen’s Magnetom 3T scanner using a 16-channel head coil with foam padding to prevent head movement. Participants were fitted with earplugs and supplied with headphones to protect hearing. During resting fMRI scans, participants were instructed to lie still with their eyes open, and to breathe and blink normally. Blood oxygen level-dependent (BOLD) data during the resting scans were recorded with a T_2_*-weighted gradient-echo echo-planar imaging sequence (EPI) and the following parameters: FOV = 220 ⨯ 220 mm, voxel size = 3.4 ⨯ 3.4 ⨯ 3.0 mm^3^, matrix size = 64 ⨯ 64, 36 interleaved axial slices per volume, TR = 2000 ms, TE = 27 ms, flip angle = 90°, GRAPPA = 2; number of volumes = 206 (scan length approx. 6.8 minutes).

DWI scans were acquired with a dual spin-echo EPI sequence and the following parameters: FOV = 256 ⨯ 256 mm, voxel size = 2.0 mm isotropic, matrix size = 128 ⨯ 128, 75 axial slices, TR = 17000 ms, TE = 98 ms, GRAPPA = 2. For each participant, two sets of DWIs were acquired with opposite phase encoding: one anterior-to-posterior (AP) and one posterior-to-anterior (PA). For each phase-encoded set, there were 80 total volumes: 60 directions with diffusion weighting factor b = 1100 s/mm^2^, 10 directions with b = 300 s/mm^2^, and 10 acquisitions with b = 0 s/mm^2^ (i.e., non-DW).

T_2_-weighted images with fat saturation were acquired with a turbo spin-echo sequence and the following parameters: FOV = 192 ⨯ 192 mm, voxel size = 1.0 ⨯ 1.0 ⨯ 1.7 mm^3^, matrix size of 192 x 192, 94 axial slices, TR = 8000 ms, TE = 89 ms, GRAPPA = 2. T_1_-weighted images were acquired with a magnetization-prepared rapid gradient echo sequence (MPRAGE) with AP phase encoding and the following parameters: FOV = 256 ⨯ 256 mm, voxel size = 1.0 mm isotropic, matrix size = 256 ⨯ 256, 176 sagittal slices, TR = 2530 ms, and TE = 3.03 ms, GRAPPA = 2.

### Resting-state preprocessing and rsFC calculations

Resting-state data were preprocessed with the AFNI software package (Cox, 1996), version 20.1.05. We describe the steps briefly here. The first 5 volumes (out of 206 acquired) were removed to ensure that magnetization was stabilized (*3dTcat)*. Preprocessing included, in order, de-spiking (*3dDespike)*, slice-timing correction to the first slice (*3dTshift*), deobliquing (*3dWarp*), motion correction (*3dvolreg*), and functional/structural affine co-registration to Talairach space using the TT_N27 anatomical template and voxel resampling to 2 mm^3^ (*align_epi_anat*.*py* and *3dAllineate*; Talairach and Tournoux, 1988). The estimates for motion correction, EPI to anatomical alignment, and transformation into standard space were combined into a single transformation so that regridding of EPI data only occurred once. Next, EPI data were spatially smoothed using a 4 mm full width at half maximum (FWHM) Gaussian kernel (*3dmerge*). We then scaled each voxel time series to a mean of 100, with a range of 0-200 (*3dTcat* and *3dcalc*; see Chen et al. 2017), regressed head motion from each voxel time series using the mean and derivatives of the 6 rigid-body parameter estimates, and detrended the data (*3dDeconvolve, 3dTproject, 3dcalc;* Power et al., 2012). Importantly, head motion correction and linear detrending were performed based on a single regression model in order to avoid the issue of reintroducing signal related to nuisance covariates caused by projecting data into a sequence of different subspaces (Lindquist et al. 2019). Prior to model regression, frames with movement displacement greater than 0.3 mm were censored. We used a threshold of 0.3 mm of average head displacement across all frames, including censored ones, during any scan to exclude participants (one participant).

After preprocessing the resting-state data, we identified the hippocampal “seed” as the voxel maximally connected with any IPC voxel for each participant, determined from the baseline fMRI scan (see *rTMS and Stimulation Localization* below). We created a 3 mm radius sphere around this location and averaged the BOLD time series of all voxels within it to derive a single hippocampal seed time series. Pearson’s R-values were then computed for the correlation between this time series and that of each voxel in the rest of the brain; these values were Fisher Z-transformed to obtain a final metric for hippocampal connectivity across the whole brain in each scan (i.e., pre- and post-stimulation). These Z-score correlation maps were used to calculate both whole-brain and hippocampal-precuneus rsFC for statistical analyses (see below).

### ROI selection for rsFC calculations

For our first analysis, we measured rsFC between the hippocampal seed, determined from the baseline resting-state scan, and a cluster of voxels in the precuneus. The boundaries of the precuneus were defined by reanalyzing the data from Wang et al. (2014). Briefly, the experimental design of Wang et al. (2014) was similar to the current work, except participants (n = 16) received both active and sham stimulation of the IPC, during separate weeks. Thus, participants underwent four scans (pre-active stimulation, post-active stimulation, pre-sham stimulation, and post-sham stimulation). First, we put the data from Wang et al. (2014) through the same preprocessing pipeline described above. Then, we performed a whole brain analysis by contrasting the changes in hippocampal rsFC following active IPC and sham stimulation. We identified a cluster of voxels spanning the left precuneus and medial occipital lobe (p < 0.05, NN = 3) and excluded voxels outside the precuneus using the CA_N27_ML AFNI atlas to form a mask of the precuneus. For the present analyses, average hippocampal rsFC was calculated in these voxels for each time point (pre- and post-stimulation) and for each participant using the AFNI *3dBrickStat* command and the correlation maps described above.

### DWI preprocessing

The combination of DW and anatomical datasets were preprocessed using a combination of FATCAT (Taylor and Saad 2013) and TORTOISE (Pierpaoli et al. 2010; Irfanoglu et al. 2018; version 3.1.4). DW DICOMs were first converted into NIFTI format (with text gradient tables; *fat_proc_convert_dcm_dwis*), and both the T_1_W and T_2_W DICOM sets were also converted into NIFTI (*fat_proc_convert_dcm_anat*). Each participant’s T_2_W image was rigidly re-oriented (“axialized”) to the standard mni_icbm152_t2 reference data set included with the FATCAT software download (*fat_proc_axialize_*anat), and each participant’s T_1_W image was then also aligned to their re-oriented T_2_W image (*fat_proc_align_anat_pair*). DW images were visually inspected, and volumes corrupted by cardiac pulsation artifacts, signal dropout, motion artifact, ghosting, and spike/radiofrequency noise artifacts were marked and removed (*fat_proc_select_vols* and *fat_proc_filter_dwis*) by authors CF or JM, and verified by MF. An average of 2.43 ± 2.35 volumes was discarded per participant (from 80 total). We then used TORTOISE’s DIFFPREP to correct for distortions caused by motion and eddy currents, in conjunction with DR-BUDDI (Irfanoglu et al. 2015) to correct for EPI distortions (B0 inhomogeneity effects). Finally, tensor fitting and tensor uncertainty estimation were performed in AFNI (*fat_proc_dwi_to_dt*), and DEC maps were generated for quality assurance (*fat_proc_decmap*). Data were inspected at multiple points of the processing pipeline for artifacts. Notably, the FATCAT and TORTOISE tools both use AFNI programs to generate images that are useful for quality control at each processing step (e.g., alignment, FA maps, etc.). Each participant’s DW image sets were inspected for artifacts before undergoing preprocessing, after preprocessing, and after tensor fitting.

### Region of interest (ROI) selection for FA calculations

FA was calculated in WM pathways between specific pairs of gray matter (GM) ROIs and restricted to the hemisphere that was stimulated (left). These GM ROIs were selected based on the theoretical mechanistic framework for rTMS-induced hippocampal network enhancement (Wang et al. 2014; Freedberg et al. 2019; Hebscher and Voss 2020). For each participant, we identified GM ROIs for the stimulation point in the IPC, parahippocampal, entorhinal, and hippocampal areas by parcellating each participant’s T_1_W scan with parcellations created by the “recon-all” command in FreeSurfer (Dale et al. 1999) version 6.0.0; in particular, we used the Desikan atlas (DK; Desikan et al. 2006) because, in general, it delineates gyral anatomy well. Note that the entorhinal cortex parcellation in this atlas corresponds to both the entorhinal and perirhinal cortices. For an example of these regions in a representative participant, see Fig. 2.

**Figure 2.**
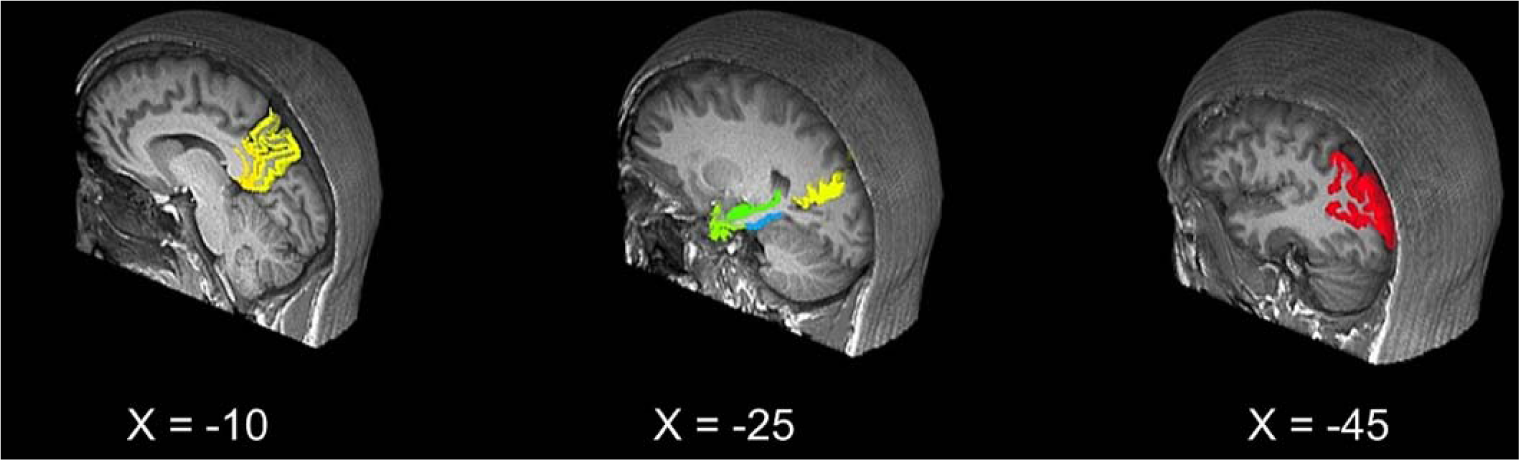
ROIs for FA calculations. ROIs are overlaid on a sample participants brain. ROIs included the precuneus (red), IPC (yellow), retrosplenial cortex (orange), parahippocampal cortex (blue), entorhinal cortex (light green), and hippocampus (dark green).

For the stimulation point GM ROI, we used the DK atlas region for the IPC, which had substantial overlap with each participant’s stimulation location. To quantify this overlap and justify our selection of the IPC to represent the stimulation location, we identified each participant’s stimulation location in Talairach space and created a 6 mm radius sphere around it. We then transformed each participant’s FreeSurfer parcellation into Talairach space and calculated the number of intersecting voxels between each participant’s sphere and each region in the DK atlas. Finally, we averaged the percentage of overlapping voxels across participants and compared the percentage of overlap between regions in the DK atlas. On average, the atlas IPC region contained the greatest density of overlapping voxels across participants (39.30%), which was substantially more than other regions, including the supramarginal gyrus (7.06%) and superior parietal cortex (1.96%). All other regions had < 1% overlap. Stimulation locations are illustrated in Fig. 3A, overlaid on a template brain. We chose the posterior cingulate cortex from the DK atlas because it corresponds well to the retrosplenial cortex (see Fig. 2).

**Figure 3.**
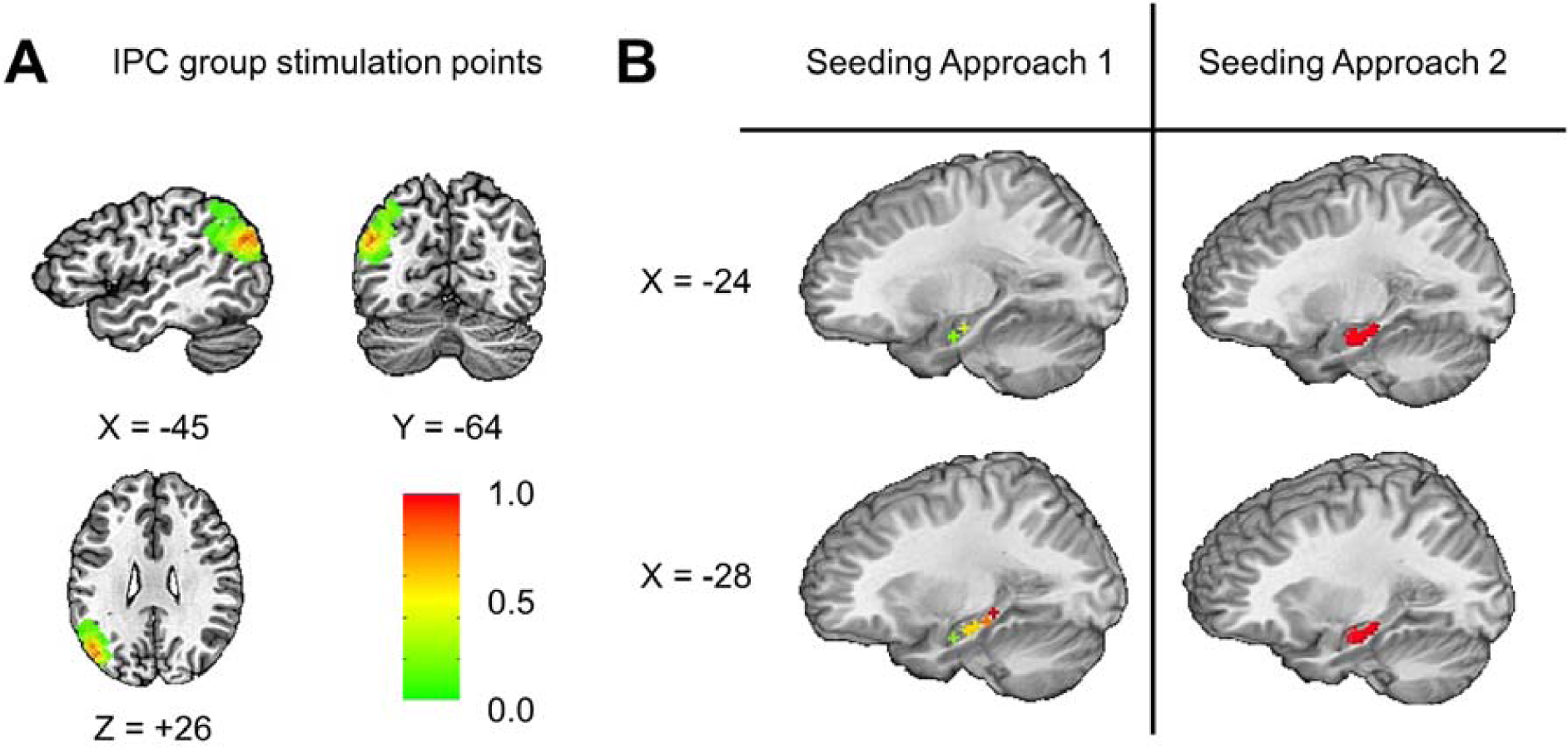
(A) Distribution of stimulation points. The topographical distribution of stimulated voxels across the left IPC are shown on a template brain (TT_N27). The color bar represents the proportion of voxels that were stimulated across participants. (B) IPC group seeding approaches. The left side shows the location of the six sampled seeds for 12 participants. The right side shows the region of voxels that were sampled to find the seed region maximally connected with the IPC.

We note that we had planned to examine the IPC-retrosplenial pathway, but were not able to produce viable tractography results in most of our participants for this pathway (see below), and thus could not analyze FA in this pathway. We also created a GM ROI of the precuneus directly from the DK atlas in each participant. Finally, we created bilateral GM ROIs in the precentral gyrus to measure FA in the corpus callosum for control analyses.

### Calculation of Fractional Anisotropy (FA)

We calculated FA in WM bundles estimated between GM ROIs using AFNI’s *3dTrackID* command (Taylor et al. 2012). These pathways were restricted to the left (stimulated) hemisphere, except for callosal fibers between precentral gyri. Prior to using *3dTrackID*, GM ROIs were inflated by up to two voxels in all directions to provide a degree of intersection with WM pathways using *3dROIMaker*, and to reduce the effects of regridding and alignment when mapping parcellations from the T_1_w dataset into the DTI grid space; this inflation was constrained by using the WM skeleton from DTI (where FA > 0.2) to limit over-inflation of ROIs into WM regions.

We performed probabilistic tractography between our GM ROIs using local uncertainties in diffusion tensor eigenvectors and the tensor-fitted data for each participant. *3dTrackID* uses repeated iterations of whole brain tracking with the FACTID algorithm (Taylor et al. 2012) to estimate the location of WM pathways between ROIs, essentially parcellating the WM skeleton into most likely pathways between ROIs based on the participant’s own diffusion data. We used a standard map where FA > 0.2 to define the WM skeleton within which to estimate tracts. The maximum acceptable “turn” angle for a pathway was set to 60 degrees. We performed 1000 iterations of whole brain tracking and all voxels through which more than five pathways passed to connect a pair of targets were included to create WM pathways associated with each pair of targets. The mean and standard deviation of the FA for each pathway were automatically calculated by the FATCAT tracking function. This resulted in a single mean FA value for each viable pathway in each participant. Outlier FA values greater or less than 3 standard deviations from the mean were disregarded.

### rTMS and Stimulation Localization

As in previous studies (e.g. Wang et al. 2014), we stimulated the sub-region of the IPC with maximum rsFC to the hippocampus in each participant. In each participant, the IPC target search volume was a sphere of 15 mm radius, cut to exclude non-brain voxels, around Talairach-Tournoux location x = −47, y = −68, z = +36 (LPI-SPM coordinate notation, here and below). We used two approaches for determining the seed and stimulus location, both employing automated scripts to remove biases in seed selection. For the first approach (12 participants), we chose the maximally connected hippocampal voxel from six pre-selected locations along the longitudinal aspect of the hippocampus in Talairach-Tournoux space (Seed 1: x = −26, y = −10, z = −17; Seed 2: x = −22, y = −16, z = −13; Seed 3: x = −30, y = −17, z = −14; Seed 4: x = −30, y = −22, z = −12; Seed 5: x = −30, y = −27, z = −9; Seed 6: x = −30, y = −32, z = − 6; Fig. 3B, left). In the second approach (12 participants), we selected the maximally connected one of 70 pre-selected voxels in the anterior hippocampus (Fig 3B, right). These included hippocampal voxels within 15 mm of the Talairach-Tournoux coordinates used by Wang et al. (2014; x = −26, y = −10, z = −17). This approach was intended to provide wider sampling within the hippocampus and to constrain the search to the part of the hippocampus related to the rTMS-induced episodic memory improvement (Wang et al. 2014).

In each approach, we created a 3 mm radius sphere around the coordinates of each voxel in the search and computed an average time series using the voxels in that sphere. We then searched the IPC sphere for the voxel with maximum correlation with the hippocampal seed, marked its location in standard space, and then back-transformed the location into participant space using the inverse matrix of the original affine transformation. Next, this location was transformed into a 3 mm radius sphere and overlaid on the participant’s structural MRI for rTMS targeting with the Brainsight frameless stereotaxic system. For the IPC target, a stimulation trajectory was created in Brainsight so that the plane of the coil was tangential to the scalp and the induced current field was oriented perpendicular to the long axis of the gyrus containing the stimulation target. For control stimulation, we located the vertex using the 10-20 International system (Steinmetz et al. 1989) and held the coil tangential to the scalp with the junction of the coil lobes in the sagittal axis (Fig. 1A).

### rTMS

TMS was delivered with a MagStim Rapid^2^ stimulator through a Double AirFilm coil. rTMS was delivered at 100% of the resting motor evoked potential threshold, which was determined immediately before the first rTMS session using the TMS Motor Threshold Assessment Tool (MTAT 2.0; http://www.clinicalresearcher.org/software.htm). Stimulation parameter settings were 2-second trains at 20-Hz (40 pulses per train) with an inter-train interval of 28 seconds. There were 40 trains, 1600 pulses, and a duration of 20 minutes per session.

### Memory Tasks

Each session of behavioral testing included an episodic memory task, a procedural memory task, a paired-associates task, and a battery of neuropsychological tests (Weintraub et al. 2013). Only the episodic and procedural memory tasks were analyzed for the current study. These two tasks were administered at the start of each behavioral testing session and their order was counterbalanced across participants to eliminate possible interfering effects (Poldrack et al. 2001; Poldrack and Packard 2003).

### Episodic memory task

The task was presented on a 22-inch monitor connected to a laptop running MATLAB (The MathWorks, Inc. Natick, MA, USA) and Psychtoolbox software (Brainard 1997; Pelli 1997; Fig. 1B, left). The task had two sections: training and testing. During training, participants viewed 20 face-word pairs. The faces were presented on the center of the screen as grayscale pictures on a white background (screen size = 2.5 x 2.5 inches) and the words were presented through speakers. The faces were taken from a database of models (Althoff and Cohen 1999) and shown for three seconds. The words were played one second after the face was presented. Testing followed training after a one-minute delay. Participants were shown each face and told to recall the word paired with each face. There was no time limit and the order of face-word pairs was different than in the training phase of the task. The proportion of successfully remembered pairs was noted after each session.

Participants saw a different set of stimuli in each behavioral session and the order of the three possible sets was counterbalanced across participants. Each set included ten male and ten female faces. Words were nouns three to eight letters long with written Kucera-Francis frequencies of 200 to 2000 and concreteness ratings of 300 to 700 (MRC Psycholinguistic Database; www.psych.rl.ac.uk).

### Procedural Memory Task

This task, based on the Weather Prediction Task (Knowlton et al. 1994), tests the ability to learn and use implicit, probabilistic, associations between events. It was performed on the same laptop computer used to administer the episodic memory task with MATLAB and Psychtoolbox software (Fig. 1B, right). Participants were told they would learn to predict one of two fictional weather outcomes (rain or sunny, hot or cold) or whether symptoms (rash, headache, sneezing, or fatigue) were associated with two fictional diseases (“nermitis” or “caldosis”), based on the presentation of four “cards” containing arbitrary stimuli (screen size = 0.9 x 1.6 inches). Each card was probabilistically associated with the outcomes based on how often it was shown and its reinforcement rate (Table 1). For each task session, each card was associated with one of the outcomes at a rate of 76, 57, 43, and 20%, as in Knowlton et al. (1994). These probabilities can be calculated for each card using the following equation and the values in Table 1:

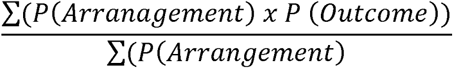

where P (Arrangement) is the probability that that card will appear on any trial, and P (Outcome) is the reinforcement rate for one of the two outcomes. Thus, for Card 1 (arrangements 8-14), the numerator was 0.343 and the denominator was 0.454, so 0.343/0.454 = 0.76, or 76%. Participants were not told these probabilities in order to discourage use of an explicit strategy.

**Table 1.** Procedural task training contingencies. Columns 2-5 from the left indicate the presence (1) or absence (0) of each card for each arrangement (Column 1). The probability that that an arrangement would appear is shown in Column 5. The reinforcement rate for each arrangement is indicated in Column 6.

| Arrangement | Card |  |  |  | P ( Arrangement) | P (Outcome) |
| --- | --- | --- | --- | --- | --- | --- |
|  | 1 | 2 | 3 | 4 |  |  |
| 1 | 0 | 0 | 0 | 1 | 0.140 | 0.15 |
| 2 | 0 | 0 | 1 | 0 | 0.084 | 0.38 |
| 3 | 0 | 0 | 1 | 1 | 0.087 | 0.10 |
| 4 | 0 | 1 | 0 | 0 | 0.084 | 0.62 |
| 5 | 0 | 1 | 0 | 1 | 0.064 | 0.18 |
| 6 | 0 | 1 | 1 | 0 | 0.047 | 0.50 |
| 7 | 0 | 1 | 1 | 1 | 0.041 | 0.21 |
| 8 | 1 | 0 | 0 | 0 | 0.140 | 0.85 |
| 9 | 1 | 0 | 0 | 1 | 0.058 | 0.50 |
| 10 | 1 | 0 | 1 | 0 | 0.064 | 0.82 |
| 11 | 1 | 1 | 0 | 1 | 0.032 | 0.43 |
| 12 | 1 | 1 | 0 | 0 | 0.087 | 0.90 |
| 13 | 1 | 1 | 0 | 1 | 0.032 | 0.57 |
| 14 | 1 | 1 | 1 | 0 | 0.041 | 0.79 |

On each trial, participants saw an arrangement of 1-3 cards and were instructed to press one of two keys on a standard keyboard to make their predictions. Correct predictions were rewarded with points. Participants started with 1,000 points and were awarded 25 points after a successful prediction or penalized by 25 points for incorrect predictions or failure to respond within 5 seconds. A prompting tone sounded two seconds after card presentation. Participants were instructed to use the strategy they felt would optimize their point winning. There were three training blocks of 50 trials, followed by a single block of 42 trials where learning was assessed in the absence of feedback.

During training, each trial began with the presentation of cards at the top of the screen. There were 14 possible unique card arrangements, which occurred with a set frequency (Table 1, P (Arrangements)). Each card was presented at a different screen position in each array to encourage participants to associate outcomes with card identities, not locations. A running point count was presented near the center of the screen in black and changed to green for successful, and red for unsuccessful, predictions. A key indicating the key-outcome mapping was shown during all trials.

Learning was calculated from the test block performance. During the test block, all aspects of the task were the same as training except that there was no time limit, the audio prompt was disabled, and the point count was replaced with question marks. Responses were scored correct when the participant chose the outcome most often associated with a successful prediction for that card arrangement during training. Participants responded to each of the 14 arrangements three times for a total of 42 trials. Since two of the arrangements predicted an outcome 50% of the time, trials with this arrangement were discarded from analysis. The task was performed in each of the three behavioral testing sessions with a different set of associations and the order was counterbalanced across participants.

### Statistical Analyses

#### Whole brain correlational analysis

To identify regions where FA was associated significantly with changes in hippocampal-cortical rsFC, we first subtracted each participant’s pre-stimulation rsFC correlation map from their post-stimulation map to form a voxel-wise correlation change map. Those maps were concatenated into a single file using *3dTcat*, where the “time” dimension was the set of participants.

We then estimated a corresponding time series of FA values of interest concatenated across participants in the same way. The corresponding “time series” of FA values was created by selecting a tractographic connection of interest that was found across all participants, and stacking the mean FA values of it. We intended to perform this analysis for three pathways of interest − 1) the IPC-parahippocampal gyrus, 2) the IPC-retrosplenial cortex, and 3) the IPC-precuneus pathways - but viable IPC-retrosplenial WM connections were only reconstructed in 83% of participants, so it was not included. As a control for an effect of individual FA as a general trait and not specific to the pathways of interest, we performed the same analysis for the IPC group using FA values for callosal connections between the left and right precentral gyri.

For each WM connection of interest, the FA “time series” was used a regressor for each voxel in the volumetric rsFC-change dataset using *3dTcorr1D*, along with a group mask to eliminate WM and ventricle voxels. This produced a single volume describing the correlation of FA and rsFC changes across the GM ribbon. Finally, we used *3drefit* to add a false discovery rate (FDR) curve to the final file.

#### Planned FA-FC correlational analysis

We regressed FA values in the IPC-parahippocampal gyrus and IPC-precuneus pathways against changes in hippocampal-precuneus rsFC for both the IPC and vertex stimulation groups (as described in the previous section), again using callosal FA as a control. All correlations were performed in R Studio (R version 3.5.1; RStudio Team, 2016) using the *cor*.*test* function. Data were checked for normality using the Shapiro-Wilk normality test, and the valid correlational test (Pearson or Spearman) was applied for the data in each case. For all correlations, one-sided tests were applied to detect significant positive correlations (i.e., increase in rsFC associated with an increase in FA), consistent with our hypotheses. The same procedures were performed for *Planned FA-Memory correlational analyses* and *Baseline rsFC and memory correlations* (see below).

#### Multivariate analysis of IPC-MTL pathways

FA from the IPC-parahippocampal, parahippocampal-entorhinal, and entorhinal-hippocampal (i.e. the MTL) pathways were fed into the *3dMVM (Chen et al. 2014)* AFNI command along with hippocampal-precuneus rsFC changes for the IPC and vertex groups. We also performed the same analysis to determine whether FA in these pathways accounted for interindividual variation in the episodic and procedural memory changes at the one and 7-14 day time points. *3dMVM* generated an omnibus F-statistic and p-value reflecting the association of all three pathways as a network, together, with our outcome measures. *3dMVM* also generates *p*-values at the level of each WM connection to determine which connections drive the network level results. For post hoc analyses, we corrected for multiple comparisons using the *fdr_bh* MATLAB command, which executes the Benjamini & Hochberg (Benjamini and Hochberg 1995) procedure for controlling the false discovery rate (FDR) of a family of hypothesis tests (Groppe 2020).

#### Planned FA-Memory correlational analyses

We regressed FA values in the IPC-parahippocampal gyrus and IPC-precuneus pathways against changes in episodic and procedural memory for the IPC and vertex stimulation groups and for the change in memory from pre-stimulation to the one and 7-14-day measurements. Again, we performed the same set of analyses for the IPC group using the callosal FA values.

#### Baseline rsFC and memory correlations

To determine whether individual variability in FA was significantly associated with baseline memory scores and rsFC changes, we repeated the above analyses but with the change scores replaced by the baseline scores for the IPC and vertex groups.

## Results

### Baseline rsFC and memory correlations

FA for the IPC-parahippocampal and IPC-precuneus pathways was not associated with baseline values in whole-brain or hippocampal-precuneus rsFC for either group (all *p* > 0.14). Pathway FA was not associated with baseline memory scores (all *p* > 0.07).

### Whole brain correlational analysis

IPC-parahippocampal FA was significantly associated with changes in hippocampal rsFC with small clusters (< 38 voxels) in the left (two clusters) and right precentral, and right inferior occipital cortex (FDR corrected, NN=3, *q* < 0.05; Table 2). Qualitatively, IPC-precuneus FA was associated with larger and more diffuse hippocampal rsFC changes compared to IPC-parahippocampal FA (FDR corrected, NN=3, *q* < 0.05; Fig. 4, right; Table 3). These regions included the left precuneus, cuneus, and fusiform gyrus. We found no voxels with hippocampal rsFC changes that were significantly related to callosal FA.

**Table 2.** Whole brain regression results for IPC-parahippocampal gyrus pathway. Each listed region represents a cluster of voxels where hippocampal-cortical rsFC was positively associated with IPC-parahippocampal gyrus FA (FDR corrected, *q* < 0.05). Coordinate notation is LPI-SPM.

| Region | Number of Voxels | Peak X | Peak Y | Peak Z |
| --- | --- | --- | --- | --- |
| Left Precentral Gyrus | 37 | -43 | -15 | +50 |
| Right Precentral Gyrus | 25 | +43 | -13 | +50 |
| Left Precentral Gyrus | 17 | -51 | -9 | +28 |
| Right Inferior Occipital Gyrus | 10 | +41 | -69 | -6 |
| Left Postcentral Gyrus | 3 | -25 | -29 | +64 |
| Left Fusiform Gyrus | 2 | -37 | -59 | -14 |
| Left Middle Occipital Gyrus | 2 | -39 | -85 | 0 |
| Right Precentral Gyrus | 2 | +55 | -5 | +36 |
| Left Paracentral Lobule | 2 | -1 | -23 | +44 |
| Left Paracentral Lobule | 2 | -5 | -33 | +60 |
| Left Postcentral Gyrus | 2 | -11 | -37 | +62 |

**Table 3.** Whole brain regression results for IPC-precuneus pathway. Each listed region represents a cluster of voxels where hippocampal-cortical rsFC were positively associated with IPC-precuneus FA (FDR corrected, *q* < 0.05). Only the top 20 largest clusters are shown. Coordinate notation is LPI-SPM.

| Region (TT_Daemon atlas) | Number of Voxels | Peak X | Peak Y | Peak Z |
| --- | --- | --- | --- | --- |
| Left Postcentral Gyrus | 2139 | -61 | -21 | +20 |
| Left Precuneus | 1981 | -1 | -55 | +46 |
| Right Precentral Gyrus | 627 | +43 | -15 | +46 |
| Right Precentral Gyrus | 552 | +57 | +1 | +6 |
| Right Fusiform Gyrus | 254 | +41 | -65 | -10 |
| Left Fusiform Gyrus | 215 | -39 | -43 | -16 |
| Right Declive (Cerebellum) | 142 | +19 | -73 | -16 |
| Right Subcallosal Gyrus | 115 | +9 | -1 | -12 |
| Left Cuneus | 102 | --3 | -83 | +28 |
| Right Superior Temporal Gyrus | 79 | +35 | +23 | -28 |
| Left Anterior Cingulate | 68 | -1 | +33 | +10 |
| Right Inferior Frontal Gyrus | 56 | +49 | +21 | +4 |
| Left Inferior Frontal Gyrus | 47 | -49 | +7 | +22 |
| Left Middle Frontal Gyrus | 47 | -21 | -13 | +60 |
| Right Inferior Temporal Gyrus | 39 | +51 | -69 | +2 |
| Left Caudate | 36 | -13 | +19 | -8 |
| Left Claustrum | 36 | -27 | +27 | 0 |
| Right Posterior Cingulate | 34 | +3 | -57 | +16 |
| Right Cingulate Gyrus | 30 | +1 | -23 | +42 |
| Left Middle Temporal Gyrus | 27 | -51 | -35 | -14 |

**Figure 4.**
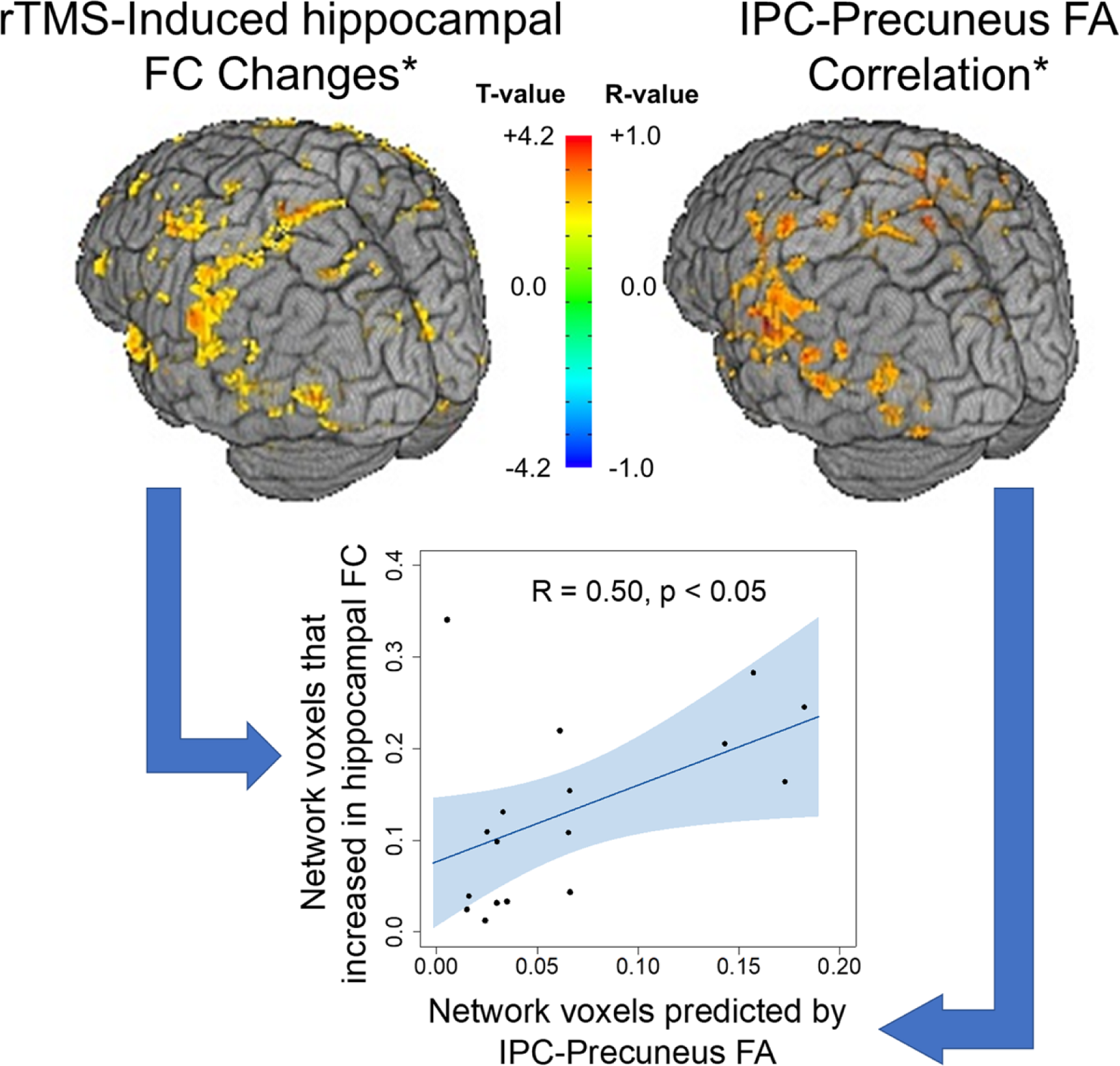
Whole brain regression results. (top left) IPC rTMS-induced changes in hippocampal rsFC from Freedberg et al. (2019). (top right) Whole brain regression examining the correlation between IPC-precuneus FA and voxel-wise changes in IPC rTMS-induced hippocampal rsFC. The color bar reflects t-values for the left image and Z-transformed R-Values for the right image.

The spatial pattern of significantly associated clusters from the IPC-precuneus whole brain analysis resembled the pattern of significant hippocampal rsFC increases found previously (Freedberg et al. 2019). To investigate this further, we separately binarized all clusters with hippocampal rsFC changes associated with IPC-precuneus FA (Fig. 4, right) and voxels with rTMS-induced hippocampal rsFC increases from Freedberg et al. (2019; Fig. 4, left). Twenty percent of the voxels that increased significantly in hippocampal rsFC following IPC stimulation were associated with IPC-precuneus FA. To test whether the distribution of regions that increased in hippocampal rsFC was similar to the distribution of hippocampal rsFC changes associated with IPC-precuneus FA, we calculated the percentage of voxels in each mask that were co-localized with 17 canonical networks (Yeo et al. 2011). Thus, for each mask, we generated 17 values representing the percentage of voxels in that mask that were in each network. Finally, we regressed the 17 values for each mask against each other. We assumed that if two masks had related topographical distributions, then the percentage of voxels from each mask in each of the 17 networks would be correlated and this was borne out (*r*(15) = 0.500, *p* < 0.05; Fig. 4, bottom). This demonstrated that the networks which exhibited significantly increased hippocampal-cortical rsFC were the same networks whose change was associated with IPC-precuneus FA. (bottom) Topographic distribution plot. Each dot represents a single network from Yeo et al. (2011). The correlation assesses the correspondence between regions where hippocampal-cortical rsFC was positively associated with IPC-precuneus FA (x-axis) and regions where IPC rTMS increased hippocampal-cortical rsFC. Axes values are represented as the proportion of voxels within each network.

### Planned FA-FC correlational analyses

For the IPC group, we found that both IPC-parahippocampal gyrus (*t*(20) = 2.68, *r* = 0.514, 95% CI [0.188 1.000], *p* < 0.001) and IPC-precuneus (*t*(22) = 2.624, *r* = 0.488m 95% CI [0.173 1.000], *p* < 0.001) FA were significantly related to the hippocampal-precuneus rsFC change (Fig. 5). Callosal FA was not related to hippocampal-precuneus rsFC change (*t*(22) = 0.749, *r* = 0.158, 95% CI [-0.197 1.000], ns). For the vertex group, neither IPC-parahippocampal (*t*(8) = 0.068, *r* = 0.024, 95% CI [-0.535 1.000], ns), nor IPC-precuneus (*t*(9) = 0.402, *r* = 0.085, 95% CI [-0.459 1.000], ns) FA were significantly related to hippocampal-precuneus rsFC change.

**Figure 5.**
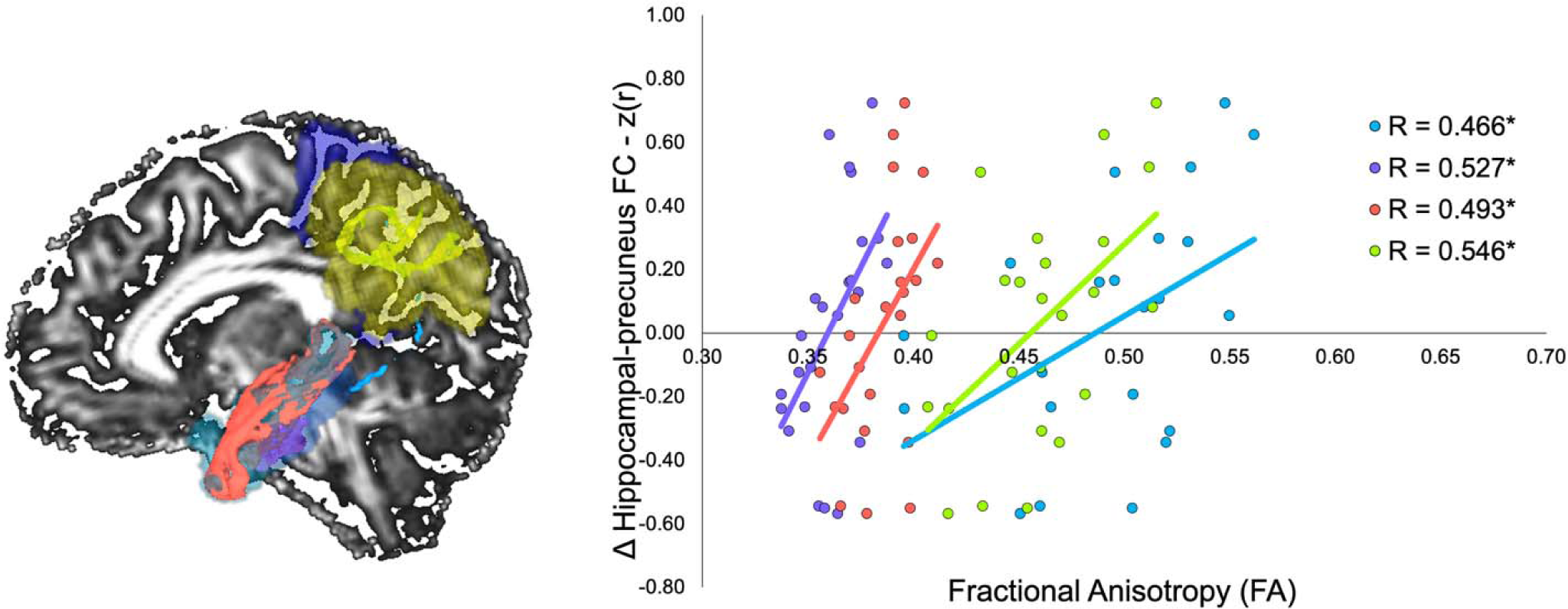
FA correlations with hippocampal-precuneus rsFC. (left) Figure shows each pathway for a sample participant, including the IPC-precuneus (yellowish-green), IPC-parahippocampal gyrus (cyan), parahippocampal gyrus-entorhinal (purple), and entorhinal-hippocampus (red) pathways. Precuneus (dark blue), IPC (yellow), and MTL (light blue) ROIs are shown as transparencies. (right) Scatter plots showing correlation between FA (x-axis) and hippocampal-precuneus rsFC. Dots and trendlines correspond to the colors of the pathways in the left figure. * - p < 0.05.

### Multivariate analysis of IPC-MTL pathways

The network level *3dMVM* model, including FA in the IPC-parahippocampal gyrus, parahippocampal gyrus-entorhinal, and entorhinal-hippocampus pathways, accounted significantly for interindividual variation in the change in hippocampal-precuneus rsFC after IPC stimulation (X^2^ = 7.801, *p* < 0.01). The Benjamini & Hochberg (1995) correction for multiple comparisons yielded a corrected *p*-value of 0.039 for the three pathway tests (IPC-parahippocampal: *t*(20) = 2.612, *p* = 0.017; parahippocampal gyrus-entorhinal: *t*(20) = 2.209, *p* = 0.039; entorhinal-hippocampal *t*(20) = 2.294, *p* = 0.033; Fig. 5). The network level *3dMVM* model for the vertex group was not significant (X^2^ = 0.678). Multivariate analyses for these pathways revealed no significant models to explain episodic or procedural memory changes (each *p* > 0.07) except for one: For the vertex group, FA between the IPC and hippocampus was negatively associated with procedural memory changes 7-14 days after stimulation (X^2^ = 11.59, *p* < 0.001). No pathways survived multiple comparisons correction.

### Planned FA-Memory correlational analyses

We regressed FA values for the IPC-parahippocampal and IPC-precuneus pathways against memory changes (Fig. 6). For the IPC group, IPC-parahippocampal FA values were correlated with episodic memory change one day after stimulation (Spearman’s correlation, *S* = 66.834, *p* < 0.001, ρ = 0.696) and at 7-14 days (*S* = 98.329, *p* < 0.05, ρ = 0.553). IPC-precuneus FA was significantly correlated with episodic memory change at one (*S* = 121.63, *p* < 0.05, ρ = 0.575) and 7-14 days after stimulation (*S* = 108.18, *p* < 0.05, ρ = 0.622). For procedural memory, no significant correlations were found for IPC-parahippocampal (one day: *t*(8) = −0.732, *r* = −0.251, 95% CI [-0.705 1.000], ns; 7-14 days: *t*(8) = −1.100, *r* = −0.362, 95% CI [-0.762 1.000], ns) or IPC-precuneus (one day: *t*(9) = 0.126, *r* = 0.042, 95% CI [-0.493 1.000], ns; 7-14 days: *t*(9) = −0.254, *r* = −0.084, 95% CI [-0.582], ns) FA. Callosal FA was not associated with episodic (one day: *S* = 207.51, ns, ρ = 0.274; 7-14 days *S* = 294.2, ns, ρ = −0.029) or procedural memory change (one day: *t*(9) = 0.706, *r* = 0.229, 95% CI [-0.335 1.000], ns; 7-14 days *t*(9) = −0.215, *r* = −0.071, 95% CI [-0.574 1.000], ns).

**Figure 6.**
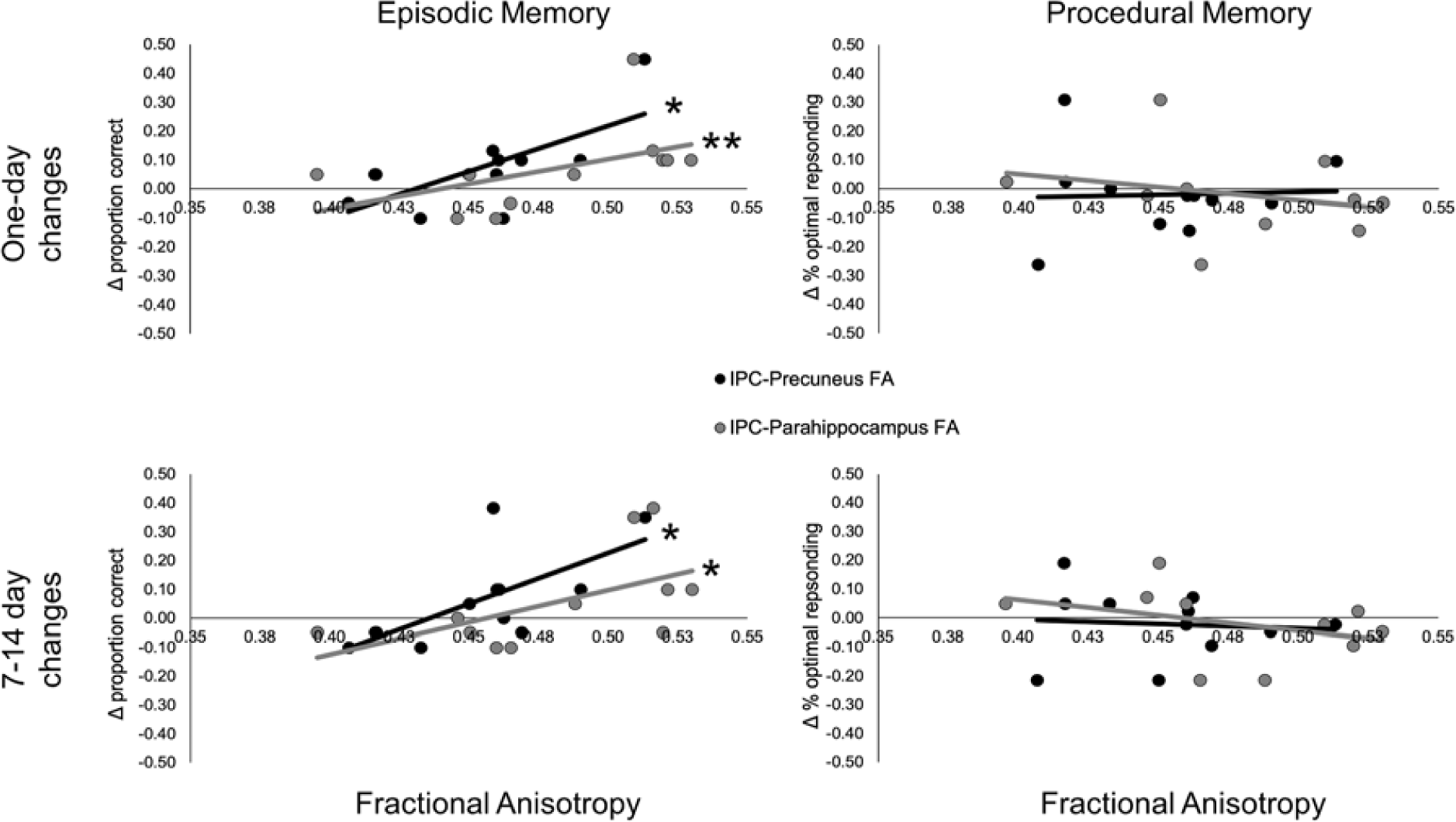
FA correlations with memory. Top plots show correlations between IPC-Precuneus (black) and IPC-parahippocampal (gray) FA (x-axes) and episodic (left plot) and procedural (right plot) memory changes one day after the last stimulation session. The bottom plots show the same data for memory changes 7-14 days after the last rTMS session. * - p < 0.05 ** < 0.001.

For the vertex group, neither IPC-parahippocampal (*S* = 191.67, ns, ρ = −0.162) nor IPC-precuneus (*S* = 232.59, ns, ρ = −0.057) FA correlated with episodic memory one day after stimulation. IPC-parahippocampal FA did not correlate with episodic memory 7-14 days after stimulation (*S* = 231.64, ns, ρ = −0.404), but IPC-precuneus FA did (*S* = 103.59, *p* < 0.05, ρ = 0.529). IPC-parahippocampal (*t*(8) = −0.335, *r* = −0.118, 95% CI [-0.693 1.000], ns) and IPC-precuneus (*t*(9) = 0.596, *r* = 0.200, 95% CI [-0.366 1.000], ns) FA did not correlate with procedural memory change after one day. We did find, 7-14 days after stimulation, that IPC-precuneus FA did correlate positively with procedural memory changes (*t*(9) = 1.896, *r* = 0.534, 95% CI [0.015 1.000], *p* < 0.05), and IPC-parahippocampal FA negatively correlated with procedural memory changes (*t*(8) = −4.351, *r* = −0.838, 95% CI [-0.961 −0.442], *p* < 0.005; two-sided test).

## Discussion

Our study links the effects of IPC rTMS to hippocampal-cortical rsFC with FA of several hippocampal network connections, supporting the hypothesis that propagation of reinforcing activity from the IPC to the rest of the network occurs through parahippocampal and precuneus WM pathways. Our results also show that while both pathways are associated with hippocampal rsFC changes, FA in the IPC-precuneus pathway correlated with widespread changes across the cortex in the same regions which were found in a previous analysis of these data to increase significantly in hippocampal rsFC following rTMS (Freedberg et al. 2019). We did not find any such effects after vertex stimulation, and there was no association of rTMS effects with callosal FA, which was our control for an effect of high FA as a general trait. FA in both pathways was associated with episodic memory changes one and 7-14 days after stimulation and no significant correlations between FA in any of these pathways and procedural memory changes after rTMS one day after stimulation. We did, however, find evidence that IPC-precuneus FA positively, and IPC-parahippocampal FA negatively, correlated with procedural memory changes 7-14 days after stimulation. These pathways may be involved in the consolidation of task-related memory, since the same correlations were not found one day after stimulation, but this will need further exploration.

The proposed mechanism for IPC rTMS-induced enhancement of hippocampal rsFC and episodic memory is propagation of signal via either IPC-parahippocampal or IPC-retrosplenial pathways (Wang et al. 2014; Freedberg et al. 2019; Hebscher and Voss 2020). Although we were unable to measure IPC-retrosplenial FA in the current work, IPC-parahippocampal FA was related to hippocampal-cortical rsFC and episodic memory changes. These results validate the hypothesized model and suggest that this pathway is critical for the effect. However, there are a few caveats. First, the multivariate model including IPC-parahippocampus, parahippocampus-enthorinal cortex, and entorhinal cortex-hippocampus FA, was not associated with episodic memory changes. Thus, it is unknown whether pathways connecting the IPC stimulation site to the hippocampus are responsible for the observed memory changes. Second, IPC-precuneus FA was associated with rsFC changes in larger, more diffuse regions across the brain, suggesting that the IPC-precuneus pathway may play a more important role in the effect of IPC rTMS on rsFC and memory. Thus, although the IPC-parahippocampus pathway is related to these effects, it may play only a complementary role with the IPC-precuneus pathway in supporting IPC rTMS effects on hippocampal rsFC and episodic memory.

While our data imply that the IPC-precuneus pathway is important for improving memory with IPC stimulation, they reveal little about its role in memory itself. Lateral parietal regions, such as the IPC and angular gyrus, and medial regions, such as the precuneus and posterior cingulate cortex, are commonly co-activated in fMRI studies of episodic memory tasks (Addis et al. 2007; Sestieri et al. 2011; Rugg and Vilberg 2013; Benoit and Schacter 2015; Ritchey and Cooper 2020), suggesting they work as a unit. Additionally, they exhibit high rsFC during rest (Greicius et al. 2003) and lower rsFC during task performance in general (Shulman et al. 1997), as do other nodes in the default mode network (DMN). The DMN is theorized to play a role in internally-directed tasks, such as episodic memory retrieval (Buckner et al. 2008). The IPC and precuneus may form a subnetwork within the DMN for episodic memory. This hypothesis is supported by the fact that these DMN regions activate early during memory recollection, as opposed to frontal regions of the DMN, such as the medial prefrontal cortex, or non-DMN regions (Sestieri et al. 2011). Periods of activity in the lateral and posterior medial cortical regions appear to demarcate the temporal boundaries of episodes identified by participants during movie-watching, suggesting that these areas play a role in segregating events during encoding (Baldassano et al. 2017). When participants in this study were read a narration of the movie, activity in the precuneus and angular gyrus showed reinstatement of event-related activity patterns just before those events were fully described, suggesting that these regions reconstruct events in real time. Thus, reinforcement of the IPC-precuneus pathway by rTMS may have facilitated the definition and reconstruction of events experienced during our episodic memory task.

The literature portrays the precuneus as involved in aspects of cognition, including visuo-spatial and mental imagery, self-referential processing, and consciousness (Cavanna and Trimble 2006), suggesting that it is important for several of the functions collected under the term “episodic memory.” The precuneus has bidirectional connections with the lateral parietal and frontal cortex, the thalamus, claustrum, striatum, and brain stem (Leichnetz 2001; Cavanna and Trimble 2006), as well as the MTL (Insausti and Muñoz 2001; Kobayashi and Amaral 2003). The fact that it has major connections with polymodal, and limited connectivity with primary, sensory regions suggests that it is involved in processing higher-order visual representations (e.g. objects, backgrounds) related to scenes or contextual information. In our task, enhancement of context could increase the binding of face-word associations. Additionally, better mental imagery may strengthen retrieval of episodic memories (e.g., word-face associations) by increasing their vividness.

Our findings are consistent with a recent meta-analysis (Beynel et al. 2020), which concluded that rTMS rsFC effects are not isolated to targeted networks or pathways. The IPC-parahippocampal gyrus and IPC-precuneus pathways both contribute to the effect of rTMS on hippocampal-cortical rsFC, even though previous studies (Wang et al. 2014; Freedberg et al. 2019) showed that IPC rTMS targeted to the sub-region with greatest hippocampal rsFC produced the greatest effect. It is also possible that the increase in hippocampal network rsFC depends on synchronous activity conducted along these two divergent pathways from the point of stimulation. This targeting uncertainty illustrates the danger of tautologically confirming posited mechanisms from behavioral outcomes and the usefulness of functional and structural imaging as adjuncts to neuromodulation.

Although we found that differences in pathway FA explained 22-30% of the variability in rTMS-induced rsFC change between individuals, there are several other possible sources of this variability (Ridding and Ziemann 2010). These include other individual factors, such as the cortical thickness of the stimulated region (Conde et al. 2012). State-specific factors that influence the effect of TMS include cortisol levels, which fluctuate throughout the day and affect plasticity (Sale et al. 2008) and menstrual cycle phase (Smith et al. 1999). We kept rTMS sessions at the same approximate time of day. However, differences in timing between participants may have also contributed to variability in response.

A design limitation of the current work is that we only examined FA in the pathways most likely to conduct the effects of IPC stimulation on hippocampal-cortical rsFC and memory, based on the anatomical and fMRI literature. Although there are several parietal (Cavada and Goldman-Rakic 1989a), prefrontal (Cavada and Goldman-Rakic 1989b), and MTL (Amaral and Witter 1989; Suzuki and Amaral 1994) regions that project to the hippocampus, we prioritized our search to avoid an excessive number of comparisons and false positive findings. A second intrinsic limitation is the limited extent to which DTI and FA can be used to characterize WM organization in voxels with crossing fibers (Alexander et al. 2001, 2002), which may exist in 63-90% of WM voxels (Jeurissen et al. 2013). High angular resolution diffusion imaging (HARDI) models can be used to improve tractography through regions of crossing fibers but requires many diffusion directions and introduces WM tracking difficulties itself (because many voxels will then contain several possible propagation pathways). We chose to acquire fewer diffusion directions in order to use the robust EPI distortion correction method in DR-BUDDI, which requires collecting DTI data in two different phase-encoding directions. This was a necessary step, as uncorrected EPI-related distortions significantly affect fiber trajectories (Irfanoglu et al. 2012). Finally, although there are properties of microstructure that determine anisotropy and conduction in WM, such as fiber density and myelination, FA is related to multiple properties of microstructure, not all of which may be related to rTMS efficacy in ways accounted for by our hypothesis.

To date, the enhancing effect of IPC rTMS on the episodic memory network and memory performance is the only finding in noninvasive neuromodulation of a cognitive system with robust and reproducible results, immediate clinical applicability and a mechanistic biomarker/surrogate endpoint whose intensity correlates with clinical response. Understanding more about its mechanism will enable further clinical and scientific advances in this area.

## Conclusion

We measured rTMS-induced changes in hippocampal rsFC and episodic and procedural memory in a sample of participants who were stimulated at the IPC region maximally connected to the hippocampus or at the vertex. DW images were collected prior to rTMS so that we could determine whether FA in pathways between the IPC and the parahippocampus and between the IPC and precuneus were associated with the effects of rTMS on rsFC and episodic memory. Our results show that while FA in both pathways were associated with rTMS-induced hippocampal-cortical rsFC changes, the association with IPC-precuneus FA encompassed larger clusters of cortex and in regions that also significantly increased in hippocampal rsFC in a prior report. FA in both pathways were also related to rTMS-induced episodic memory changes. These effects were not found when substituting FA values in pathways unrelated to the effect of stimulation in the group that received IPC rTMS. Further, for participants who received stimulation of the IPC, we did not observe an association between FA and rTMS-induced effects on procedural memory, which relies on a different network than the one targeted. These results suggest that both the IPC-parahippocampal and IPC-precuneus pathways play a role in the enhancing effect of stimulation on hippocampal rsFC and episodic memory.

## Acknowledgments

We would like to thank Dr. Joel Voss at Northwestern University for sharing his data for this study.

